# Cross-Protection Against Zika Virus Infection Conferred by a Live Attenuated Japanese Encephalitis SA14-14-2 Vaccine

**DOI:** 10.1101/2020.02.14.950352

**Authors:** Ran Wang, Zida Zhen, Lance Turtle, Baohua Hou, Yueqi Li, Na Gao, Dongying Fan, Hui Chen, Jing An

**Author notes:** Corresponding author: Hui Chen,.

## Abstract

Zika virus (ZIKV) and Japanese encephalitis virus (JEV) are closely related mosquito-borne flaviviruses. Japanese encephalitis (JE) vaccine SA14-14-2 has been in the Chinese national Expanded Program on Immunization since 2007. The recent recognition of severe disease syndromes associated with ZIKV, and the identification of ZIKV from mosquitoes in China, prompts an urgent need to investigate the potential interaction between the two. In this study, we showed that SA14-14-2 is protective against ZIKV infection in mice. JE vaccine SA14-14-2 triggered both Th1 and Th2 cross-reactive immune responses to ZIKV; however, it was cellular immunity that predominantly mediated cross-protection against ZIKV infection. Passive transfer of immune sera did not result in significant cross-protection, but did mediate antibody dependent enhancement *in vitro*, though this did not have an adverse impact on survival. This study suggests that SA14-14-2 vaccine can protect against ZIKV through a cross-reactive T cell response. This is vital information in terms of ZIKV prevention or precaution in those ZIKV-affected regions where JEV circulates or SA14-14-2 is in widespread use, and opens a promising avenue into developing a novel bivalent vaccine against both ZIKV and JEV.

**Importance:** Japanese encephalitis is a controllable disease in many countries in Asia, especially in China, where many people have Japanese encephalitis virus (JEV) immunity due to extensive JEV vaccination campaigns or natural exposure. Live-attenuated SA14-14-2 strain is a safe and effective vaccine recommended by the World Health Organization and has been vaccinated more than 600 million doses since 1989. As the prevalence of Zika virus (ZIKV) and rising risk in above regions, the cross-reactive immune response between these two antigenically closely related flaviviruses, JEV and ZIKV, should also be fully recognized, which is presumed to be based on those ambiguous cross-reactive immunity between dengue virus and ZIKV. In this study, we found that JEV SA14-14-2 vaccine conferred cross-protection against ZIKV challenge in mice, which is mainly due to cellular immunity rather than neutralizing antibody response. However, specific protective components or cooperation between components warrant to be explored in subsequent experiments. In conclusion, this study can provide important evidence for those who live in JEV-endemic areas and are at risk for ZIKV infection.

## Introduction

Recently, Zika virus (ZIKV) has caused devastating outbreaks of fetal congenital malformations in South and Central America and now transmitted in more than 70 countries, including many previously unaffected regions. ZIKV infection during pregnancy increases the risk of neurological disorders in newborns (1), such as microcephaly. In adults, ZIKV causes Guillain-Barré syndrome and other neurologic disorders (2). So far, no specific vaccine or antiviral for prevention and treatment of Zika have been licensed.

ZIKV is a member of the genus *Flavivirus*, family *Flaviviridae*, which contains more than 70 viruses. Among of them, mosquito-borne flaviviruses such as Japanese encephalitis (JE) virus (JEV), dengue virus (DENV), ZIKV, yellow fever virus, and West Nile virus pose a threat to half of the world population and cause significant public health impact in many developing countries (3).

The flavivirus genome consists of non-segmented single-stranded positive-sense RNA, which encodes three structural proteins including the capsid protein (C), the membrane protein (M) and the envelope protein (E), and seven non-structural (NS) proteins (NS1, NS2a, NS2b, NS3, NS4a, NS4b, and NS5). Within the same genus, these mosquito-borne flaviviruses are antigenically related to various degrees (4). Among them, JEV, ZIKV and DENV share more than 50% amino acid sequence identity by pairs (5, 6). On average, ZIKV shares a 55.6% protein sequence identity with DENV and 56.1% with JEV (5).

Currently, several studies have indicated complex interactions between DENV and ZIKV immunity (7-12). Clinical data suggest that pre-existing DENV immunity is partially protective against symptomatic ZIKV infection and against congenital ZIKV syndrome (13, 14). Previous DENV infection also probably partially protects against JE indicating the possibility of a more general effect within the genus (15).

Previously, we demonstrated that JEV SA14-14-4 live attenuated vaccine, an inactivated vaccine based on P3 strain, and a JE DNA vaccine based on the premembrane and E proteins, effectively elicited the production of cross-reactive antibodies, cytokines, and cellular immune responses, and generated cross-protection against four serotypes of DENV (16, 17). However, little is known about cross-reactivity between JEV and ZIKV. The geographic overlap, possibility of sequential infection with JEV and ZIKV, and widespread use of JEV SA14-14-2 vaccine in China indicate a need to understand the impact of pre-existing immunity to JEV (acquired either through SA14-14-2 vaccination or natural infection) on ZIKV infection. Therefore, in this study we aimed to evaluate the cross-reactivity and cross-protection of JEV SA-14-14-2 vaccination against ZIKV infection in a mouse model. Our findings suggest cross-immunity among between JE vaccine and ZIKV, and indicate a need for further study in humans to address the role of JE immunity in protection from ZIKV.

## Materials and Methods

### Ethical approval

The animal experiments were performed in strict accordance with the recommendations in the national guidelines for the use of animals in scientific research “Regulations for the Administration of Affairs Concerning Experimental Animals”. All experimental procedures were approved by the Institutional Animal Care and Use Committee of Capital Medical University, China. All efforts were made to minimize suffering.

### Cells, viruses, vaccine and mice

C6/36 cells were cultured in RPMI1640 medium (Gibco, USA) supplemented with 10% fetal bovine serum (FBS, Gibco, USA) at 28 °C. Vero cells were cultured in minimal essential medium (MEM, Gibco, USA) supplemented with 5% FBS at 37 °C. THP-1 cells were cultured in RPMI1640 medium supplemented with 10% FBS at 37 °C. All cells were cultivated under a humidified atmosphere of 5% CO_2_.

The JEV (Beijing-1 strain) and the ZIKV (SMGC-1 strain) were propagated in C6/36 cell cultures and stored at –80 °C. Virus titers were determined by plaque assay on Vero cells under MEM with 1.2% methylcellulose overlay medium. Virus was harvested infected C6/36 cell supernatant, concentrated by 8% polyethylene glycol precipitation, and purified from clarified extracts by ultracentrifugation. The JE live-attenuated vaccine (SA14-14-2 strain) was produced by the Chengdu Institute of Biological Products (China).

Female and male C57BL/6 mice, and female *Ifnar*^-/-^ mice were housed under specific-pathogen-free conditions, and C57BL/6 mice were bred to obtain neonatal mice. Adult female mice were used at six weeks of age, and neonatal *Ifnar*^-/-^ mice were used between 24 h and 36 h after birth.

### Mouse immunization

Six-week-old female adult mice were divided randomly into vaccine and control groups. C57BL/6 and *Ifnar*^-/-^ mice in the vaccine group were immunized intraperitoneally (i.p.) with 10^4^ and 10^3^ plaque forming units (PFU) JEV SA14-14-2 strain, respectively, three times at three-week intervals (16). Control mice were injected with PBS following an identical schedule.

### Cross-reactive protection against ZIKV in SA14-14-2-immunized *Ifnar*^-/-^ mice

Three weeks after the final vaccination, at 15 weeks of age, the *Ifnar*^-/-^ mice were challenged i.p. with a lethal dose of JEV or ZIKV (10^3^ PFU for both). Body weight and mortality were monitored daily for 14 consecutive days.

### Plaque reduction neutralization test (PRNT)

Sera were collected from the C57BL/6 mice three weeks after the final vaccination. Neutralizing antibody (nAb) titers were detected by measuring plaque reduction neutralization titer (PRNT) as previously reported (18, 19). Serum samples were heated at 56 °C for 30 minutes to inactivate complement and then two-fold serially diluted from 1:10 to 1:1,280. Diluted sera were mixed 1:1 with virus suspension containing 50 plaque-forming units (PFU), and incubated at 37 °C for 1 h. The mixture was transferred to a confluent monolayer of Vero cells in a 24-well plate, and incubated at 37 °C for another 1 h. After washing, the infected Vero cells were overlaid with MEM containing 1.2% methylcellulose followed by incubation at 37 °C for five–eight days. Plaques were visualized by crystal violet counter staining and counted. The reciprocal highest serum dilution yielding a 50% reduction of the average number of plaques as compared with the virus infection wells was calculated as the 50% neutralization titer (PRNT_50_).

### *In vitro* neutralizing and passive cross-protective effects of immune sera in neonatal C57BL/6 mice

Sera were collected from C57BL/6 mice three weeks after the final vaccination. After heat inactivation, pooled sera were mixed with either live JEV (Beijing-1 strain) or the ZIKV (SMGC-1 strain) followed by incubation at 37 °C for 1 h. Subsequently, 10 μl of serum/virus mixture containing 150 PFU of virus was gently injected intracerebally (i.c.) into neonatal C57BL/6 mice. The mice were monitored daily for body weight and mortality for 31 consecutive days.

### Enzyme-linked immunosorbent assay (ELISA)

To detect IgG antibodies and their subclasses, ELISA was performed according to the method as previously described (18). Sera were collected from C57BL/6 mice three weeks after the final vaccination. Each well of 96-well plates was coated with purified viral particles (10^5^ PFU) from JEV or ZIKV and then blocked with 3% bovine serum albumin. Sera from immunized mice were two-fold serially diluted in PBS (from 1:100 to 1:204,800), IgG antibodies were measured with goat-anti-mouse secondary antibodies (1:4,000, alkaline phosphatase coupled, Abcam, USA) and substrate solution of *p*-nitrophenyl phosphate (Sigma, USA). The optical density (O.D.) at 405 nm was measured using an ELISA reader (Thermo, USA). The reciprocal of the highest dilution, that yielded the O.D. value greater than half of the O.D. value of corresponding control at 1:100 dilution, was recorded as the end-point titer of IgG antibody (18).

To determine IgG subclasses, mouse sera (at 1:100 dilution) were used as the primary antibodies, IgG1, IgG2a, IgG2b and IgG3 subsets were detected with goat-anti-mouse secondary antibodies (1:4,000, alkaline phosphatase coupled, Abcam, USA) and substrate solution of *p*-nitrophenyl phosphate, and the levels of IgG subclasses were recorded as O.D. value at 405 nm.

### Antibody-dependent enhancement (ADE) assay

Three weeks after the final immunization, sera were collected from C57BL/6 mice. Serially ten-fold-diluted sera were incubated with JEV or ZIKV at a multiplicity of infection of 2 for one hour at 37 °C before adding to THP-1 cells. Samples were incubated for two hours at 37 °C and gently shaken every 15 min. Cells were centrifuged and washed three times, followed by resuspension in fresh RPMI-1640 medium and incubated for three days at 37 °C. Supernatants were collected and viral titer measured by plaque assay on Vero cells as described (18). Fold enhancement was calculated by comparison to viral titers in the absence of immune sera.

### Enzyme-linked immunospot (ELISPOT) assays

Three weeks after the final immunization, the cytokines IL-2, IL-4, and IFN-γ secreted by splenocytes of C57BL/6 mice were determined using ELISPOT kits (BD, USA) according to the manufacturer’s instructions. In brief, mouse splenocytes were plated at 3 × 10^5^ / well into 96-well filtration plates (Millipore, USA) pre-coated with capture antibodies and stimulated with 5 µg / well purified JEV or ZIKV particles for 60 h at 37 °C. After incubation with biotinylated detection antibody, the spots were visualized by adding streptavidin-horseradish peroxidase and 3-amino-9-ethylcarbazole substrate, counted automatically with an ELISPOT reader (CTL, USA) and analyzed by ImmunoSpot software (version 5.1). Splenocytes cocultured with concanavalin A served as a positive control, and those cultured with RPMI 1640 medium served as a negative control.

### Adoptive transfer and cross-reactive protection of splenocytes from SA14-14-2-immunized C57BL/6 mice in *Ifnar*^-/-^ mice

Three weeks after the final vaccination, splenocytes were isolated from the immunized C57BL/6 mice using lymphocyte separation percoll (Solarbio, China). A total of 3 × 10^6^ lymphocytes was transfused retro-orbitally (r.o.) to female adult *Ifnar*^*-/-*^ mice. After 24 h, the mice were challenged i.c. with a lethal dose of either JEV (Beijing-1 strain, 10^4^ PFU) or ZIKV (SMGC-1 strain, 10^4^ PFU). Body weight and mortality were monitored daily for 14 consecutive days. Mice exhibiting more than 25% loss in body weight were humanely euthanized for ethical reasons.

### Statistical analysis

Statistical analyses were performed using SPSS statistics version 17.0 (SPSS Inc., USA). Geometric mean titers (GMTs) were calculated after log transformation of reciprocal titers. Weight changes were analyzed by repeated measures analysis of variance. Kaplan-Meier survival curves were plotted and evaluated statistically by Log-rank test. Others were analyzed using one-way analysis of variance. The results were presented as means + / - / ± standard deviation (SD), and the difference between means is considered significant if *P* < 0.05. *P* values are denoted (*) if *P* < 0.05, (**) if *P* < 0.01, and (***) if *P* < 0.001, respectively.

## Results

### Cross-protection against lethal ZIKV challenge in immunocompromised mice

A number of studies have used *Ifnar*^*-/-*^ mice to evaluate the effectiveness of vaccines despite the immunodeficiency of the mice (20, 21). Although *Ifnar*^*-/-*^ mice lack the innate type 1 IFN response, they retain adaptive immunity but are highly susceptible to ZIKV (22), representing a suitable model for testing vaccine induced cross-protective adaptive immunity. First, we used this model to determine whether JEV SA14-14-2 has a cross-protective effect on ZIKV. Three doses of JE vaccine SA14-14-2 were administrated to *Ifnar*^*-/-*^ mice, followed by challenge with JEV or ZIKV three weeks post the final immunization (Fig. 1A). As expected, after JEV challenge, all mice inoculated with JEV SA14-14-2 survived (6/6) during the observation period without any weight loss (Fig. 1B and 1C). In contrast, the control mice showed a rapid decrease one day post JEV challenge and all mice died by the humane endpoint on day 3 post challenge. Interestingly, SA14-14-2-vaccinated *Ifnar*^*-/-*^ mice were completely protected (6/6) against ZIKV infection, as compared with PBS-treated mice where 83.3% (5/6) mice succumbed to infection (Fig. 1B and 1C). Vaccinated mice maintained a normal body weight whereas control mice showed significant weight loss (approximately 16%) and most of them died within 9 days (1/6 survival). These data suggest that vaccination with SA14-14-2 in *Ifnar*^*-/-*^ mice induced *in vivo* cross-reactive immunity, conferring effective cross-protection against lethal ZIKV infection.

**FIG 1.**
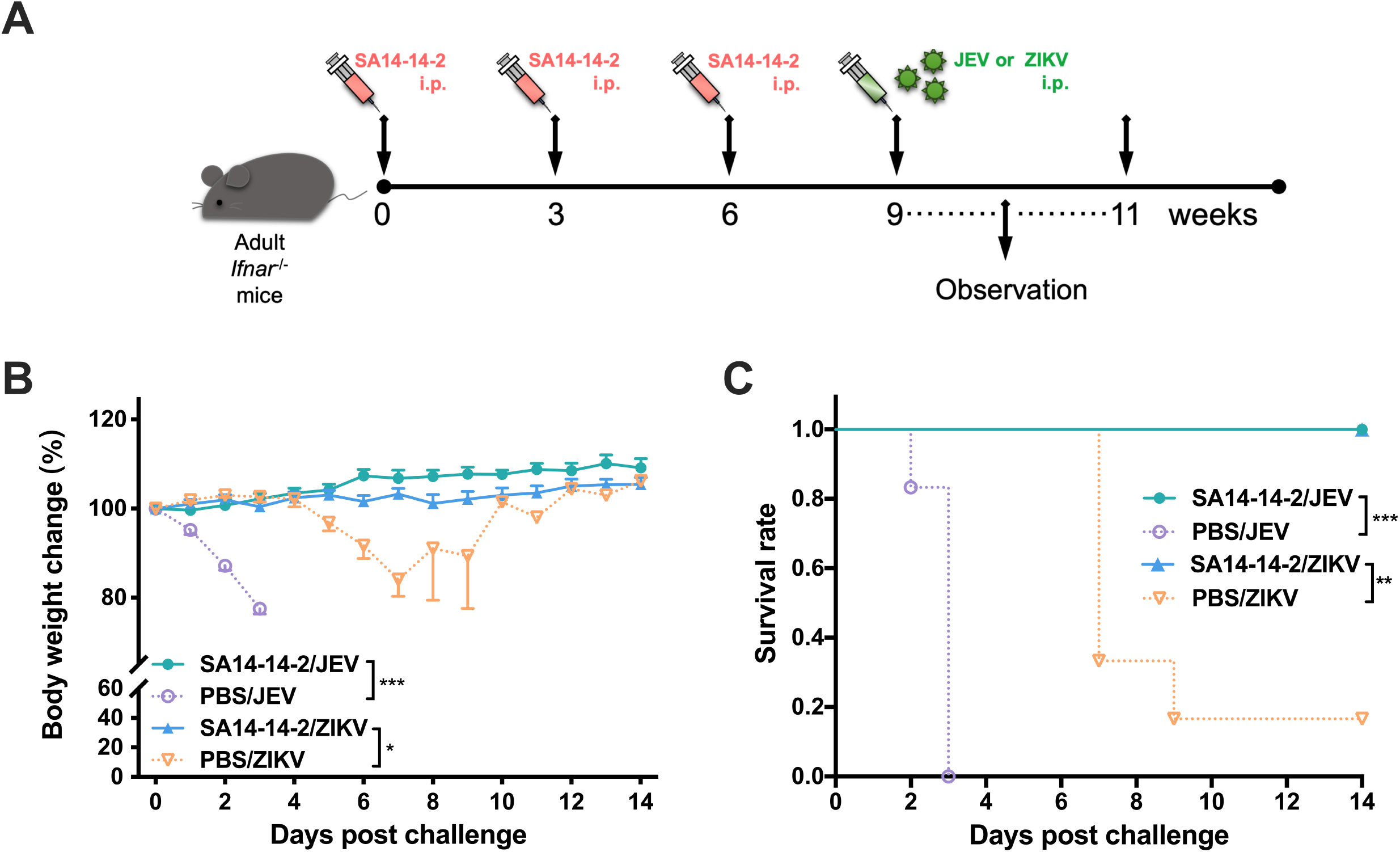
Cross-reactive protection against ZIKV in SA14-14-2-immunized *Ifnar*^-/-^ mice. (A) Schedule. Female *Ifnar*^-/-^ mice were immunized three times at three-week intervals. Three weeks after the final vaccination, the *Ifnar*^-/-^ mice were challenged i.c. with a lethal dose of JEV or ZIKV. Body weight and mortality were monitored daily for 14 consecutive days. (B) Percentage changes of body weight. Data were expressed as mean +/- SD. (C) Survival rate was shown as the percentage of survivors (*n* = 6). Mice exhibiting more than 25% loss in weight were humanely euthanized for ethical reasons. Each experiment was independently repeated three times. \**P* < 0.05; \*\**P* < 0.01; \*\*\**P* < 0.001.

### Cross-protection against ZIKV infection is not mediated by SA14-14-2-immune sera

There is a clearly established role for nAb in protection from flaviviruses. Therefore, in order to evaluate cross-reactive neutralization of ZIKV induced by SA14-14-2 vaccination, three weeks after the final immunization, sera were collected from C57BL/6 mice and the nAb titers were assayed by PRNT (Fig. 2A). As expected, mice administered three doses of JEV SA14-14-2 developed a high level of JEV-specific nAb, with a GMT of 1:494 (Fig. 2B). In contrast, JEV SA14-14-2 antisera from the immunized mice failed to neutralize ZIKV, and the PRNT_50_ titer of antisera was comparable to that of controls. This result suggests that nAb elicited by JEV SA14-14-2 yielded no neutralizing activity against ZIKV.

To further test whether JEV SA14-14-2 antisera would cross-protect mice against ZIKV challenge *in vitro*, for example by a mechanism distinct from neutralization, immune sera were harvested from SA14-14-2-vaccinated C57BL/6 mice on week three after the final immunization. A serum/virus (JEV or ZIKV) mixture was prepared and inoculated i.c. into naïve neonatal C57BL/6 mice (Fig. 2A). Control sera from C57BL/6 mice injected with PBS were included in the same experiment. SA14-14-2-immune serum protected neonatal mice (8/8) from JEV challenge, after which the mice exhbited steady growth and development, manifesting a normal and continuous weight increase, reaching 15.4 g at the end of observation (Fig. 2C and D). In contrast, none of the neonatal mice (0/8) receiving non-immune sera survived JEV challenge, and exhibited severe growth delay, with an endpoint weight of only 2.0 g. ZIKV challenge was less pathogenic in this model than JEV, with control mice (ZIKV infected receiving no sera) developing subnormally but with indistinctive body weight change (Fig. 2C). No effect of SA14-14-2-immune serum could be detected in this experiment, the terminal weights of the mice receiving JEV SA14-14-2-immune serum and control mice were 9.8 g and 9.1 g, respectively. Survival in the two groups was also the same, 10.0% (1/10) in mice incoculated with mixture of SA14-14-2-immune sera and ZIKV and 8.3% in mice injected with control mixture (1/12, Fig. 2C and D). Although SA14-14-2-immune sera did slightly extend the median survival of neonatal ZIKV challenged mice (22.0 days *vs*. 15.0 days), this effect was not significant by Log-rank test. These data suggest a potent JEV-specific protective effect of SA14-14-2-immune sera, but no cross-protection from ZIKV infection, consistent with the nAb data.

**FIG 2.**
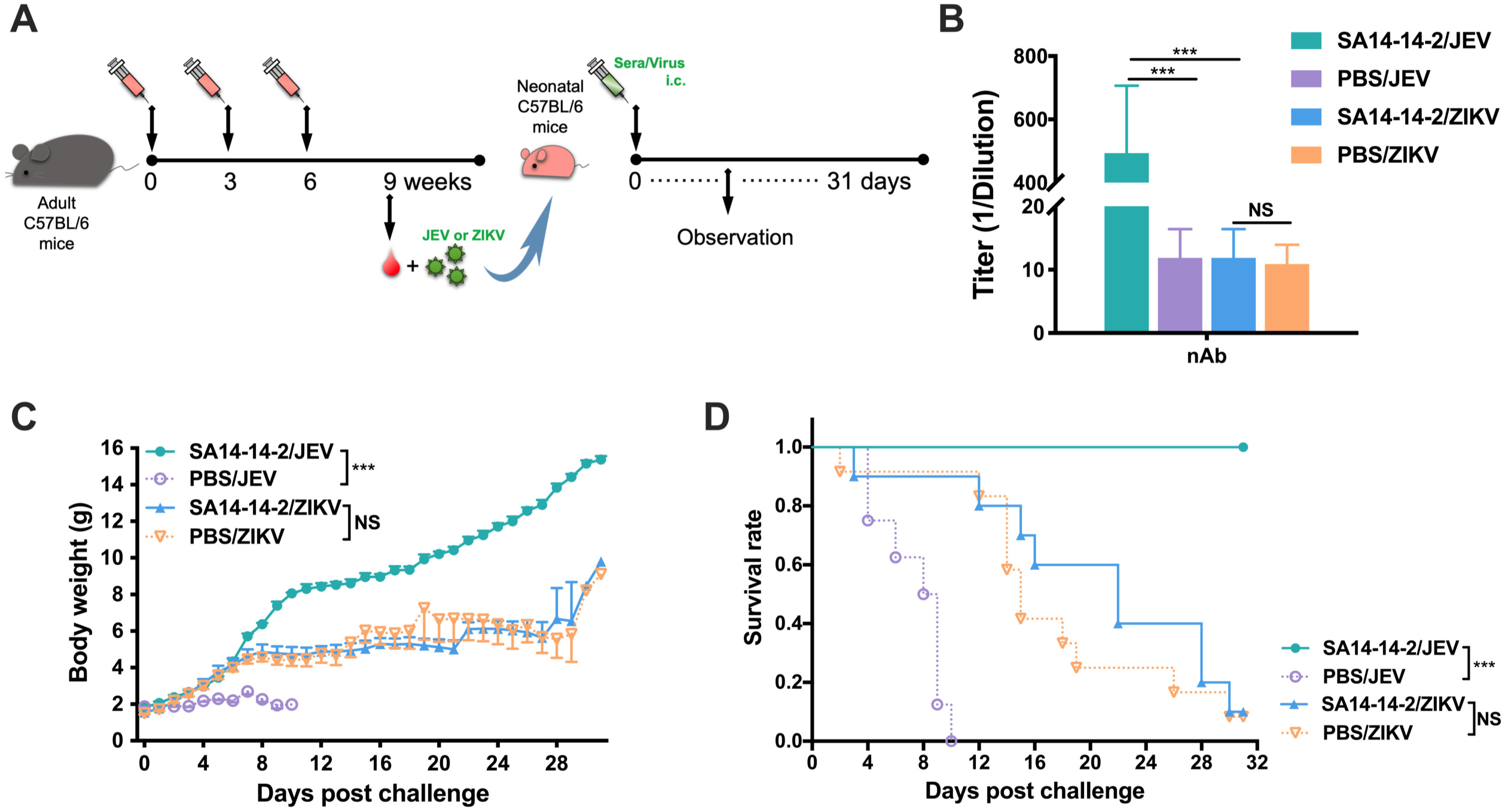
Cross-reactive nAb responses in mouse sera. (A) Schedule. Female adult C57BL/6 mice were immunized as previously described. Sera were collected and mixed with either JEV (Beijing-1 strain) or ZIKV (SMGC-1 strain). Then, serum/virus mixture was gently injected i.c. into neonatal C57BL/6 mice. (B) Serum cross-reactive nAb titers assayed by PRNT_50_ (*n* = 7). NAb titers are expressed as GMT + SD. (C) and (D) *In vitro* neutralizing and passive cross-protective effects of immune sera (*n* = 8, 8, 10, 12, respectively). The mice were monitored daily for body weight and survival rate for 31 consecutive days. (C) Body weight was expressed as mean +/- SD. (D) Survival rate was shown as the percentage of survivors. \*\*\**P* < 0.001; NS, non-significant.

### Presence of cross-reactive IgG antibody and its multiple subclasses in response to ZIKV induced by SA14-14-2-vaccination

Whilst we could detect no protective role of cross-reactive antibody against ZIKV infection from SA14-14-2-immune sera, we sought to determine whether there was any binding to ZIKV, as cross-reactive, binding but non-neutralizing antibodies have been described (23). Thus, to determine the presence of JEV-specific and ZIKV-cross-reactive IgG antibody and its subclasses, including IgG1, IgG2a, IgG2b, and IgG3, induced by SA14-14-2-vaccination, sera were collected three weeks after the last immunization from C57BL/6 mice and analyzed by ELISA (Fig. 3A). Levels of both JEV-specific and ZIKV-cross-reactive IgG antibodies were higher in the sera of immunized mice than those in corresponding controls (1:60,887 *vs*. 1:519 and 1:1,345 *vs*. 1:436 endpoint titers, respectively, Fig. 3B), suggesting the induction of a cross-reactive humoral immune response to ZIKV. Furthermore, to informatively characterize the profile of cross-reactive IgG antibodies induced by SA14-14-2-vaccination, IgG1, IgG2a, IgG2b, and IgG3 subclasses in immune sera were measured (Fig. 3C). It is known that isotype switching to IgG1 is promoted by a Th2 response, whereas switching to IgG2a, IgG2b, and IgG3 is promoted by a Th1 response (24). Our data show that IgG1, IgG2a, and IgG2b were all detected after vaccination, but IgG3 was not induced. These data suggest that SA14-14-2-vaccination could induce a response with both Th1 and Th2 components.

**FIG 3.**
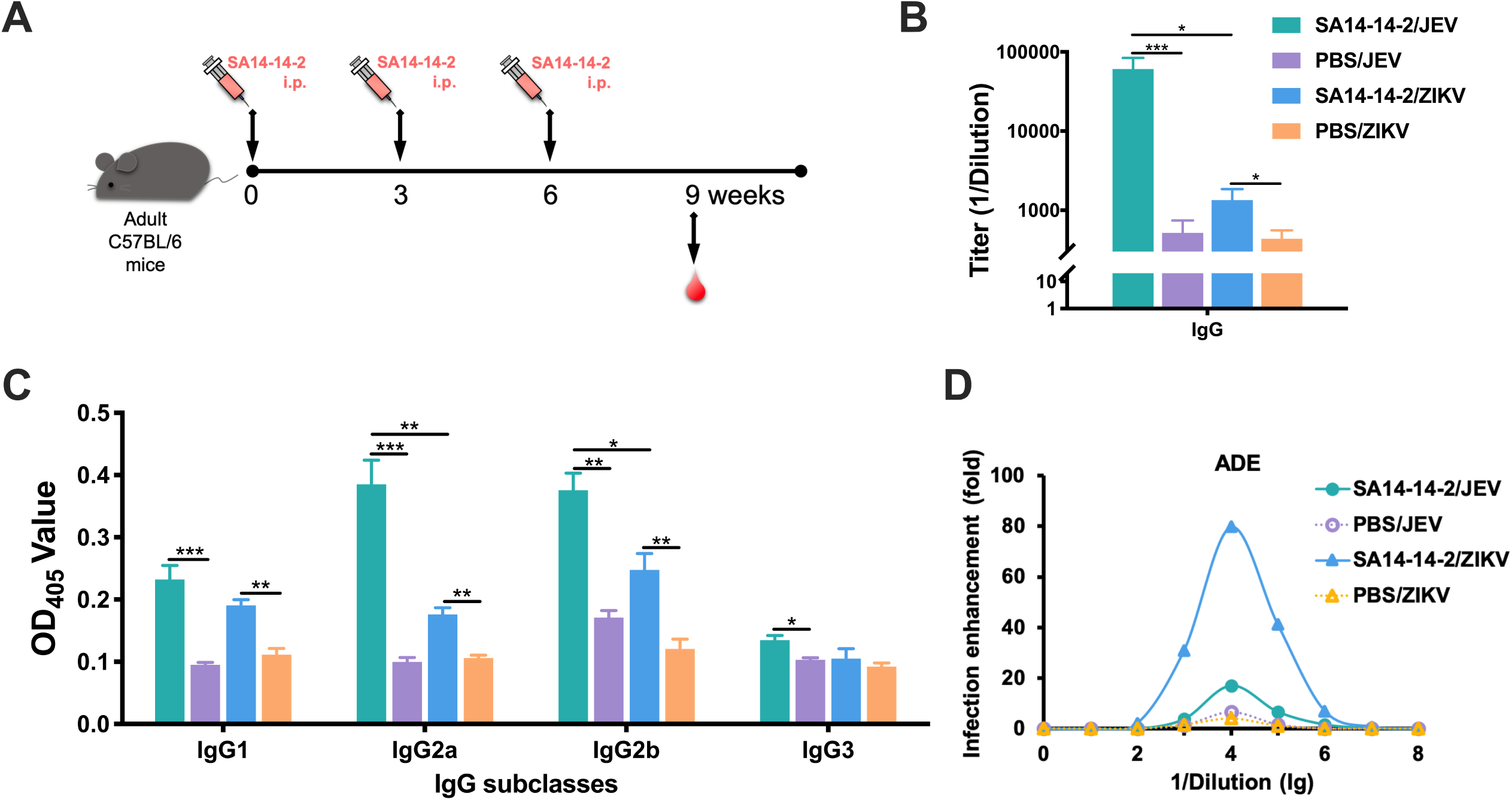
Cross-reactive IgG and its subclass responses and ADE in mouse sera. (A) Schedule of mouse immunization and serum collection. Female adult C57BL/6 mice were immunized three times at three-week intervals. Sera were collected three weeks after the final immunization. (B) Cross-reactive IgG responses detected by ELISA (*n* = 7). Ab titers are recorded as GMT + SD. (C) Cross-reactive IgG subclass responses determined by ELISA (*n* = 7). (D) ADE in sera measured by plaque forming (*n* = 7 per dilution). \**P* < 0.05; \*\**P* < 0.01; \*\*\**P* < 0.001.

### SA14-14-2-immune sera induce antibody dependent enhancement (ADE) *in vitro*

ADE occurs when the titer or neutralization potential of serum is too low to achieve complete neutralization, but the antibody is still able to bind to the virus, which then promotes entry into Fc receptor bearing cells, which are permissive for viral replication. The effect is common among flaviviruses (25). Therefore, in order to determine whether SA14-14-2-immune sera could promote ADE of ZIKV infectivity *in vitro*, we exposed FcγRI/II-bearing cell line THP-1 to ZIKV in the presence or absence of SA14-14-2-immune sera. We observed dose dependent enhancement of infection from a dilution of 1:100, which peaked at 1:10,000 dilution, with ZIKV infection enhancement up to 79.7-fold (Fig. 3D). In contrast, sera from control mice did not significantly enhance the infectivity of ZIKV, although modest enhancement at a dilution of 1:10,000, likely to due to non-specific effect. Meanwhile, when THP-1 cells were infected in the presence of the SA14-14-2-immune sera, we found that they also yielded a 17.0-fold greater infection of JEV at a dilution of 1:10,000 than those in the absence of sera (Fig. 3D). This homotypical enhancement is most likely the result of sub- or non-neutralizing titer of serum dilution. As expected, control sera did not obviously enhance JEV infection. In summary, the result demonstrates that cross-reactive anti-JEV antibodies can promote ADE of ZIKV, at least *in vitro*.

### Multiple cross-reactive cytokine responses to ZIKV

Having provided indirect evidence that both Th1 and Th2 responses were made following SA14-14-2 vaccination, and that these responses could cross-react with ZIKV, albeit not with protective nAb, we sought to determine the nature of the T helper response after SA14-14-2-vaccination. Three weeks after the third and final immunization, splenocytes were collected from the C57BL/6 mice. The levels of splenocyte-derived IL-2, IL-4, and IFN-γ were determined by ELISPOT assay (Fig. 4). Notably, when pulsed with ZIKV antigen, splenocytes from SA-14-14-2-immunized mice responded by making all three cytokines, although the levels were lower when compared with responses of SA-14-14-2-immune splenocytes upon stimulation with JEV antigen. IL-2 and IFN-γ are predominant markers of the Th1 response, IL-4 expression is defined as a marker of the Th2 response. The results indicated that either the Th1- or the Th2-type cross-reactive immune responses against ZIKV was evoked by administration of the JEV SA-14-14-2 vaccine.

**FIG 4.**
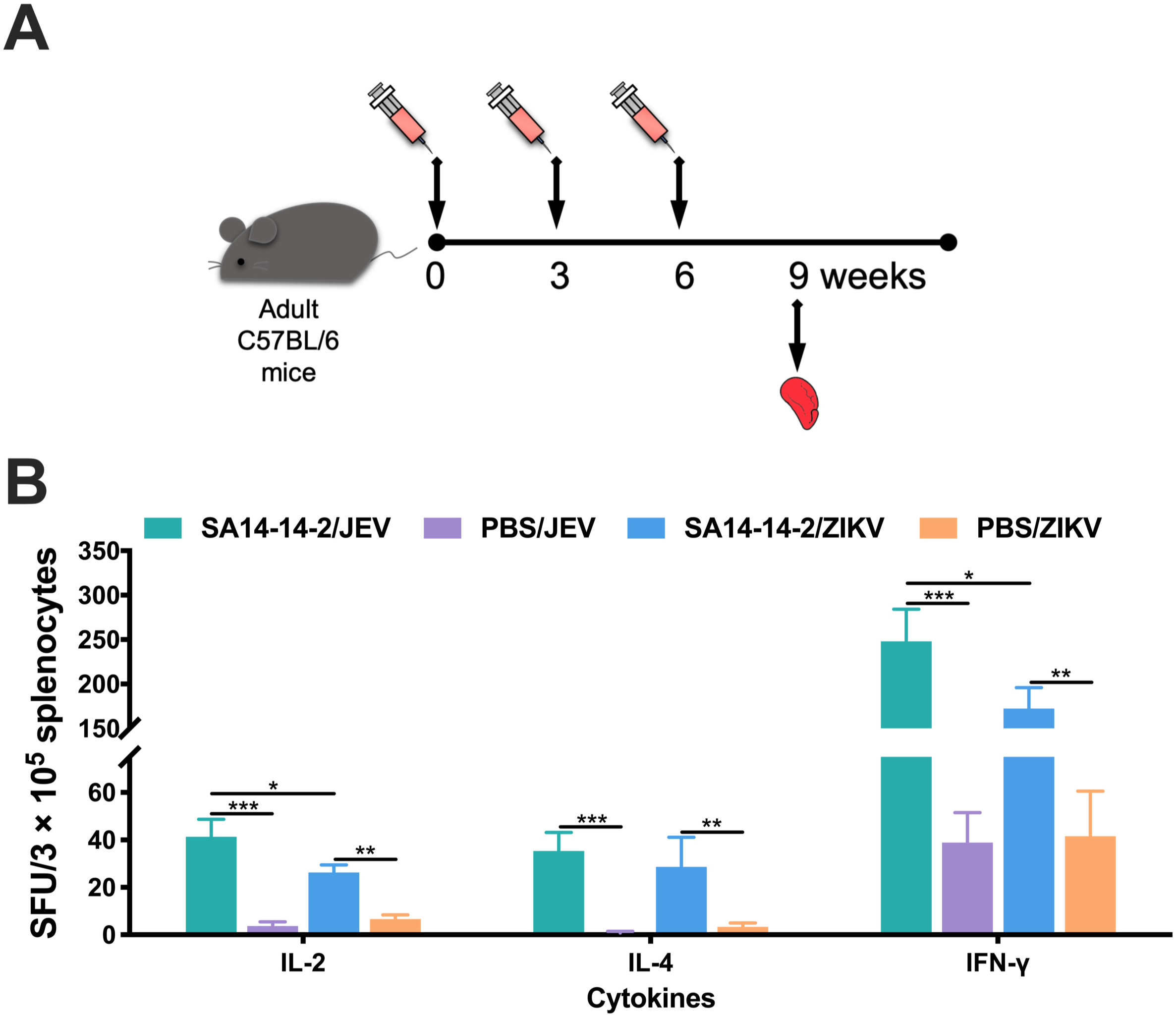
Cross-reactive cytokine responses in mouse splenocytes. (A) Schedule. Three weeks after the final immunization, splenocytes from adult C57BL/6 mice were collected. (B) The cytokines IL-2, IL-4, and IFN-γ secreted by splenocytes were determined using ELISPOT (*n* = 7). The numbers of cytokine-positive cells are reported as the mean SFU / 3 × 10^5^ splenocytes + SD. \**P* < 0.05; \*\**P* < 0.01; \*\*\**P* < 0.001.

### Cell-mediated immunity as a potential mechanism of cross-protection against ZIKV

Although multiple cytokines can be produced under ZIKV antigen stimulation, it was unclear whether cellular immunity was indispensible or essential for the *in vivo* cross-reactive protection. Therefore, adoptive transfer of immune splenocytes from SA14-14-2-immunized C57BL/6 mice into naïve *Ifnar*^*-/-*^ recipient mice was performed three weeks after the final immunization, followed by viral challenge with a lethal dose of JEV or ZIKV (Fig. 5A). After JEV challenge, mice receiving splenocytes derived from control group showed marked and up to 26.7% body weight loss, all mice died (0/6) within nine days, whereas 100% (6/6) of mice receiving SA14-14-2-immune splenic lymphocytes survived without obvious weight change (Fig. 5B and C), suggesting that a protective prototype of splenocytes activated by SA14-14-2 was successfully established. Meanwhile, while naïve *Ifnar*^*-/-*^ recipient mice in control group showed high susceptibility to ZIKV infection with weight loss of 17.0% and only 16.7% (1/6) survival rate, virtually 83.3% (5/6) of mice infused with SA14-14-2-immune lymphocytes survived the lethal ZIKV challenge with only a weight loss of 5.8% (Fig. 5B and C). Taken together with our nAb data, this result demonstrated that a cell-mediated cross-reactive response induced by SA14-14-2-immunization was protective against subsequent ZIKV infection.

**FIG 5.**
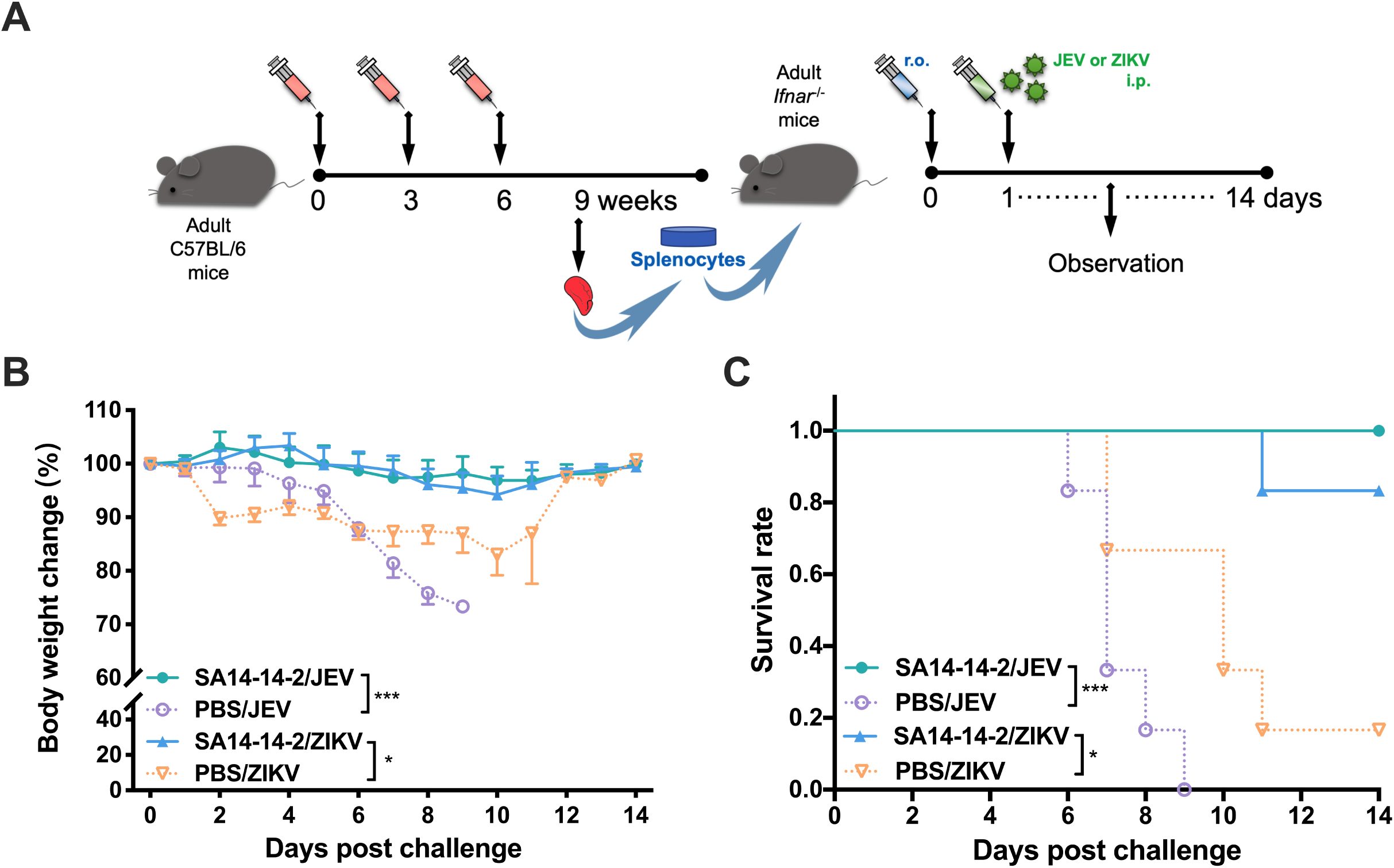
Adoptive transfer of splenocytes from SA14-14-2-vaccinated C57BL/6 mice and cross-protection of splenocytes in *Ifnar*^-/-^ mice. (A) Schedule. Splenic lymphocytes (3 × 10^6^ cells per mouse) collected three weeks post final immunization were adoptively transferred r.o. to naïve adult *Ifnar*^*-/-*^ mice, and one day later mice were challenged with a lethal dose of either JEV or ZIKV. As a control, splenic lymphocytes from PBS-treated mice were transferred to naïve *Ifnar*^*-/-*^ mice prior to challenge. Results were evaluated for (B) body weight change and (C) survival rate of mice 14 consecutive days post challenge (*n* = 6). Mice exhibiting more than 25% loss in weight were humanely euthanized for ethical reasons. Each experiment was independently repeated three times. Results are expressed as mean +/- SD. \**P* < 0.05; \*\*\**P* < 0.001.

## Discussion

It is becoming increasingly apparent that the pre-existing immunity triggered by primary flavivirus infection or vaccination can induce immunological cross-reactivity to secondary exposure with a genetically and antigenically closely related flavivirus. Immunological cross-reactions have been implicated in both protection and pathology, and there is some controversy on the consequences to the outcome of infection. An *in vitro* study showed enhancement of ZIKV replication in the presence of DENV antibodies (23). However, clinical cohort and case control studies of individuals in dengue endemic regions suggest the opposite, that pre-existing dengue immunity reduces the risk of symptomatic ZIKV infection and congenital ZIKV syndrome (13, 14, 26). Pantoja *et al*. studied the effects of pre-existing DENV immunity on ZIKV infection *in vivo* in rhesus macaques, and confirmed that the previous exposure to DENV didn’t result in enhancement of ZIKV pathogenesis (7). Intriguingly, there has been relatively little ZIKV infection reported in Asia, and it has been suggested that cross-reactive T cell responses generated by *Culex*-borne flaviviruses, of which the principle agent is JEV, has limited ZIKV spread in Asia (27).

Previously, we found that live attenuated JE vaccine SA-14-14-2 conferred cross-reactive nAbs which contributed to the cross-protection against DENV challenge (16). In contrast to this result, in this study, mice immunized with SA14-14-2 showed a cross-reactive IgG antibody response to ZIKV without the presence of neutralizing activity (Figs. 2B and 3B). Although the ADE in SA14-14-2-immune sera was detected *in vitro* (Fig. 3D), we found no evidence that this resulted in a harmful effect; indeed, in subsequent ZIKV challenge, SA14-14-2 vaccination slightly lengthened the median survival (Fig. 2D). This is an important result, suggesting that cross-reactive antibodies from SA14-14-2 vaccination are not pathogenic *in vivo*, because many people in Asia have received inactivated JE vaccine which will generate anti-JEV nAb but may not contain many of the cross-reactive T cell epitopes, which lie in the NS proteins (28). Here, we use neonatal C57BL/6 mice instead of neonatal *Ifnar*^*-/-*^ mice because the former is susceptible to ZIKV and can mimic the signs (18). Although SA14-14-2 vaccine triggered both Th1 and Th2 responses (Figs. 4B and 3C), adoptive splenocyte transfer was superior to serum transfer in protection against ZIKV infection, implying a strong correlation between SA14-14-2-induced cellular immunity and the ZIKV-cross-protective capacity. However, the limitation in this study is that we did not elucidate the functional components which are cross-protective by further sorting of splenocytes for adoptive transfer, such as purified T cells or their subpopulations (CD8^+^ or CD4^+^ alone), or cross-reactive memory B cells. Furthermore, we did not determine whether the cross-reactive nAb response was present in recipient mice transferred with SA14-14-2-immune splenocytes, although this is unlikely to change our primary conclusions.

Traditionally, based on cross-neutralizing activity, flaviviruses have been subdivided into distinct serocomplexes. Cross neutralization between different serocomplexes is usually not observed (4). The flavivirus E protein is the principal antigen against which the nAb response is directed. The extent of cross-neutralization correlates with the amino acid sequence identity of E protein: when the sequence identity in E protein is less than 40%, cross neutralization is lost (29). JEV and ZIKV belong to distinct serocomplexes, although the homology of the E protein amino acid sequence between the JEV SA14-14-2 strain and the ZIKV SMGC-1 strain was 53.4%, the distinct epitopes in E proteins of JEV and ZIKV within different flaviviruses that dominate antibody responses are presumably responsible for the unavailable cross-neutralization (30). For the sequence homology of the E protein, ZIKV is more closely related to the DENV than to the JEV serocomplex (4). However, most B cell epitopes are conformational (31), and therefore, sequence homology may not fully reflect “relatedness” as measured by cross-reactive humoral responses. Interestingly, one structural model of the relatedness of flavivirus surface topology in fact placed ZIKV and JEV closer to each other than either was to DENV (32). Nevertheless, despite this structural similarity, this did not account for protection in the model we describe here.

In contrast, NS proteins among flaviviruses are more conserved with up to 68.0% identity than structural proteins. Weiskopf *et al*. indicated that, following heterologous DENV infection, memory CD8^+^ T cells expanded that recognized conserved NS proteins (33). In fact, NS3 and NS5 represent the main targets of the CD8^+^ T cell response to flaviviruses (34, 35). One study reported more cross-reactive T cell responses to full length NS3 helicase, because of higher sequence homology, than that to the protease region alone (36). Also, a homologous analysis based on the NS5 protein would place ZIKV closer to the JEV serocomplex than to DENVs (37). In our study, we hypothesise that it is the cross-reactive response to shared T cell epitopes in the NS proteins that contributes to protection mediated by adoptive transfer of immune splenocytes.

In a mouse model, CD8^+^ T cells can mediate protection against ZIKV (35). DENV-specific cross-reactive CD8^+^ T cells can also protect (38), and similar responses are detected in humans, where cross-reactive CD8^+^ T cells specific for DENV display anti-ZIKV effector potential toward ZIKV, mediating direct cytolysis (39). JEV / ZIKV share 63.9% and 68.0% homology in NS3 and NS5, respectively. Our findings are consistent with a cross-reactive CD8^+^ T cell mediating protection, but characterization of the key components responsible for cross-protection in splenic lymphocytes, and identification of JEV/ZIKV cross-reactive epitopes, warrant further investigation.

Since 2007, China has included a two-shot schedule of the JEV SA14-14-2 vaccine into the national Expanded Program on Immunization. Children under the age of 13 and even older children in some provinces of China have pre-existing JEV immunity. It should be noted that, according to the eatablished schedule, children are received the SA14-14-2 vaccine at eight months and two years of age, but three immunizations in mice were performed in this study, as previously (16). In order to ensure comparability with what humans receive, future studies may need to test various dosing regimens.

Fu *et al*. and Xiao *et al*. identified ZIKV in mosquitoes in Guizhou province and Yunnan province (40, 41), indicating that ZIKV is present in China. Therefore, it is inevitable that JEV and ZIKV co-circulate. The identification of ZIKV presents new challenges for prevention and control because of its severe consequences in pregnancy (42), and prompts an urgent need to clarify the mechanisms underlying such cross-protective immunity, which will inform the strategy of developing safe and effective vaccines targeting both viruses, with appropriately balanced B cell and T cell antigens.

In conclusion, the results of our study demonstrated that JEV SA14-14-2 elicited effective cross-protection against ZIKV in mice. Our results indicate the potential for the widespread use of the vaccine, especially in those co-circulating countries. Moreover, this study will provide important information in terms of ZIKV prevention or precaution. Furthermore, it is worthwhile to identify common epitopes for the future development of a novel bivalent vaccine based on T cell against both JEV and ZIKV.

## Acknowledgments

This study was supported by the National Natural Science Foundation of China (grant numbers 81772172 and 81671971), and the Research Project of Beijing Children’s Hospital, Capital Medical University, National Center for Children’s Health, China (grant number GPQN201909), which funded the experimental work. Lance Turtle is supported by the Wellcome Trust (grant number 205228/Z/16/Z) and EU Horizon 2020 ZikaPLAN (Preparedness Latin America Network) consortium (grant agreement No. 734584).

R.W. performed experiments, analyzed data, and wrote the manuscript, Z.Z., B.H., Y.L. helped to perform experiments, L.T. interpreted data, and assisted in writing the manuscript, N.G. and D.F. provided valuable suggestions for manuscripts and experiments, H.C. designed the experiment and reviewed the manuscript, J.A. reviewed and guided the overall experiment. All authors have critically read and edited the manuscript.

